# microRNA-mediated regulation of microRNA machinery controls cell fate decisions

**DOI:** 10.1101/2021.07.21.453175

**Authors:** Qiuying Liu, Mariah K. Novak, Rachel Pepin, Taylor Eich, Wenqian Hu

## Abstract

microRNAs associate with Argonaute proteins, forming the microRNA-induced silencing complex (miRISC), to repress target gene expression post-transcriptionally. Although microRNAs are critical regulators in mammalian cell differentiation, our understanding of how microRNA machinery, such as the miRISC, are regulated during development is still limited. We previously showed that repressing the production of one Argonaute protein, Ago2, by Trim71 is important for mouse embryonic stem cells (mESC) self-renewal (Liu et al., 2021). Here we show that among the four Argonaute proteins in mammals, Ago2 is the major developmentally regulated Argonaute protein in mESCs. Moreover, in pluripotency, besides the Trim71-mediated regulation of *Ago2* (Liu *et al*., 2021), microRNA-182/microRNA-183 also repress *Ago2*. Specific inhibition of this microRNA-mediated repression results in stemness defects and accelerated differentiation through the let-7 microRNA pathway. These results reveal a microRNA-mediated regulatory circuit on microRNA machinery that is critical to maintaining pluripotency.

## Introduction

microRNAs (miRNAs) are endogenous ∼22-nucleotide(nt) RNAs with critical roles in modulating gene expression under diverse biological contexts (Bartel, 2009; 2018). Most miRNAs are produced from long primary transcripts (pri-miRNAs) through successive processing by two double-strand RNA (dsRNA) endonucleases named Drosha and Dicer, generating pre-miRNAs and ∼22-nt dsRNAs, respectively. One RNA strand in the ∼22-nt dsRNA, the mature miRNA, is selectively incorporated into the Argonaute (Ago) protein, forming the miRNA-induced silencing complex (miRISC) (Ha and Kim, 2014). In animals, miRISC recognizes its target mRNAs through partial base-pairings mediated by the miRNA (Bartel, 2009). The Ago protein recruits GW182 proteins to down-regulate target mRNA expression through mRNA degradation and/or translational repression (Nilsen, 2007). Although microRNAs play critical regulatory roles in mammalian cell differentiation (Ameres and Zamore, 2013; Ebert and Sharp, 2012), our understanding on how microRNA machinery, particularly the miRISC, are regulated during development is still limited.

We recently found that Ago2, a key component in the miRISC, is repressed at the mRNA translation level by an RNA-binding protein named Trim71 in mouse embryonic stem cells (mESCs) (Liu *et al*., 2021). This repression of *Ago2* inhibits stem cell differentiation mediated by the conserved pro-differentiation let-7 miRNAs (Bussing et al., 2008; Liu *et al*., 2021). These results suggest that *Ago2* is developmentally regulated during stem cell self-renewal and differentiation, and beg for characterization of additional regulators of *Ago2*. Moreover, besides Ago2, there are three additional Ago proteins (Ago1, Ago3, Ago4) in mammals that function redundantly in the miRNA pathway (Meister, 2013). The relative abundance of these Ago proteins and their contribution to miRNA activities during cell differentiation, however, are still unknown.

Here, using mESC fate decisions between pluripotency and differentiation as a mammalian cell differentiation model, we determined that Ago2 is the predominant Ago protein in mESCs, and Ago2 level increases when mESCs exit pluripotency. In the pluripotent state, miR-182 and miR-183, two conserved miRNAs abundantly expressed in mESCs, repress *Ago2* and control the stemness of mESCs. Specific inhibition of miR-182/miR-183-mediated repression of *Ago2* results in stemness defects and accelerated differentiation of mESCs through the let-7 microRNA pathway. Collectively, these results reveal a miRNA-mediated regulatory circuit on the miRNA machinery that is critical to maintaining pluripotency.

## Results

### Ago2 is the predominant developmentally regulated Argonaute protein in mESCs

Mammals have four Ago proteins (Ago1-4) that function redundantly in miRNA-mediated regulations (Meister, 2013). Transcriptomic profiling on mESCs from different laboratories indicated that mESCs express only *Ago1* and *Ago2* (Figure 1 – Figure supplement 1A) (Liu *et al*., 2021; Marks et al., 2012). To examine the relative abundance of Ago1 and Ago2 at the protein level, we generated mESCs with a Flag-tag knocked-in at the N-terminus of the Ago1 and Ago2 loci, respectively, via CRISPR/Cas9-mediated genome-editing (Figure 1 – Figure supplement 1B&C). These mESCs with the Flag-tag knocked-in displayed no stemness defects compared to the WT mESCs (Figure 1 – Figure supplement 1D) and enabled us to use the same antibody (e.g., anti-Flag) to compare the relative abundance of Ago1 and Ago2. Western blotting via an anti-Flag antibody indicated that Ago2 is the predominant Ago protein in mESCs at the protein level (Figure 1A).

**Figure 1.**
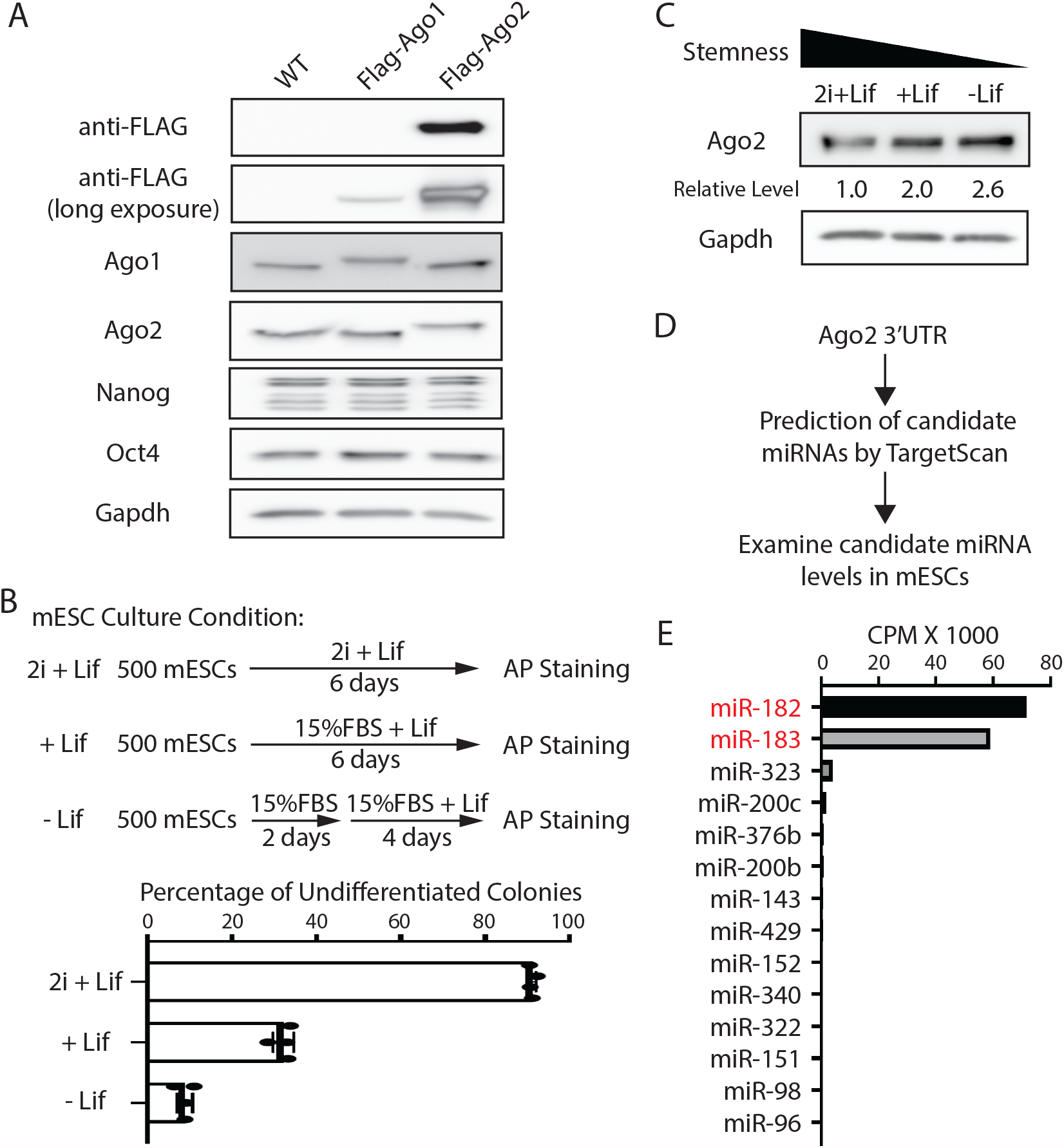
Ago2 is the major developmentally regulated Argonaute protein in mESCs. A. Western blotting in the WT, Flag-Ago1, and Flag-Ago2 mESCs. B. Colony formation assay for the mESCs. The WT mESCs were cultured under the indicated conditions, and the resultant colonies were fixed and stained for AP (alkaline phosphatase activity). The results represent the means (± SD) of four independent experiments. C. Western blotting in the WT mESCs cultured under the indicated conditions. D. Outline of identifying miRNAs that can potentially regulate *Ago2*. E. Expression levels of the identified miRNAs from (D) in mESCs. CPM: counts per million reads. The following figure supplements are available for Figure 1: Figure supplement 1. Expression of Argonaute proteins in mESCs. Figure supplement 2. miR-182 and miR-183 are associated with *Ago2* mRNA in mESCs.

To examine whether Ago2 level is regulated during mESCs differentiation, we cultured mESCs under three different conditions that mimic three different developmental stages: ground/naive state (in 2i+Lif), primed state (in 15%FBS+Lif), and differentiating state (in 15%FBS without Lif), which resulted in decreasing stemness in mESCs, as determined by the colony formation assay (Figure 1B). Western blotting indicated that Ago2 level increased when mESCs exited pluripotency (Figure 1C). This result indicated that Ago2 is developmentally regulated in mESCs, and Ago2 level is repressed in the pluripotent state.

### miR-182/miR-183 regulate *Ago2* and maintain stemness in mESCs

To determine how *Ago2* is regulated in mESCs, we hypothesized that miRNAs expressed in mESCs might contribute to the repression of *Ago2* because miRNAs are important negative regulators of gene expression. We identified the conserved miRNA-binding sites in the 3’UTR of *Ago2* mRNA through TargetScan (Agarwal et al., 2015) and then examined the expression level of the corresponding miRNAs in mESCs using existing small-RNA-seq datasets (Liu *et al*., 2021) (Figure 1D). This analysis revealed that among the miRNAs that can potentially regulate *Ago2*, miR-182 and miR-183, two miRNAs from the same miRNA family that are abundantly expressed in stem cells (Dambal et al., 2015), have significantly higher expression levels (Figure 1E).

Interestingly, miRNA-182/miR-183 decrease when mESCs transition from the ground state to the primed and differentiating state (Hadjimichael et al., 2016; Wang et al., 2017), which negatively correlates with the Ago2 expression pattern during this transition (Figure 1C). These observations suggest that *Ago2* is repressed by miR-182/miR-183 in mESCs. Consistent with this notion, using RNA antisense purification, we found that miR-182 and miRNA-183 specifically associated with *Ago2* mRNA in mESCs (Figure 1 – Figure supplement 2).

Two lines of evidence indicated that miR-182/miR-183 regulate *Ago2* mRNA. First, Ago2 increased when *miR-182, miR-183*, or both *miR-182* and *miR-183* were knocked out in mESCs (Figure 2 – Figure supplement 1A & Figure 2A&B). Second, when either miR-182 or miR-183 was over-expressed in the wild-type (WT) mESCs (Figure 2 – Figure supplement 1B), the Ago2 level decreased (Figure 2 – Figure supplement 1C). The results from these loss-of-function and gain-of-function experiments argue that miR-182/miR-183 repress *Ago2* expression in mESCs.

**Figure 2.**
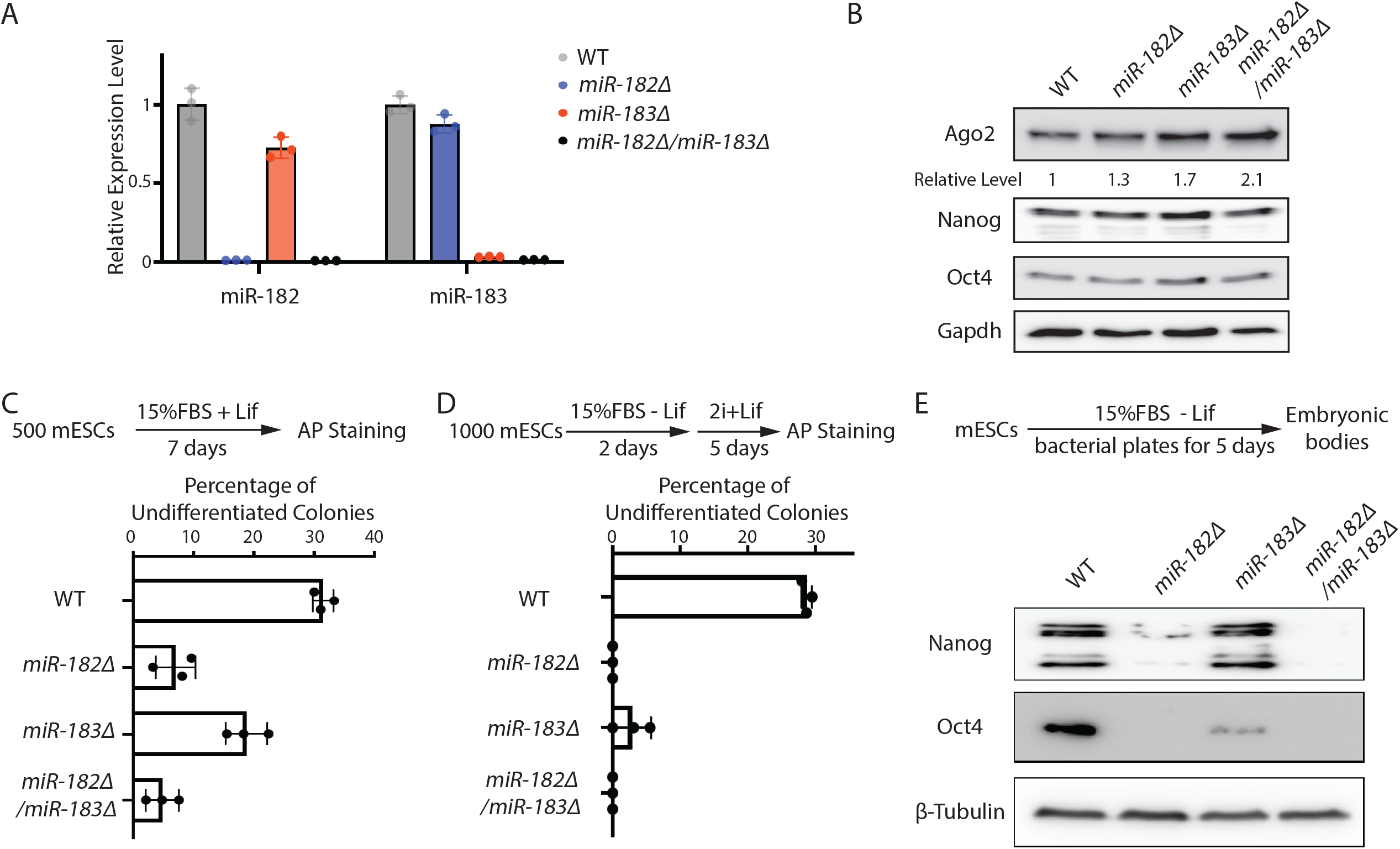
miR-182/miR-183 regulate *Ago2* and maintain stemness in mESCs. A. qRT-PCR on miR-182 and miR-183. For each miRNA, the expression level in WT cells was set as 1 for relative comparison. U6 RNA was used for normalization. The results represent the means (± SD) of three independent replicates. B. Western blotting in the WT, *miR-182Δ*, *miR-183Δ*, and *miR-182Δ/ miR-183Δ* mESCs. Gapdh was used for normalization in calculating the relative expression levels. C. Colony formation assay for mESCs. The mESCs were cultured in 15%FBS + Lif for 7 days, and the resultant colonies were fixed and stained for AP. D. Exit pluripotency assay for mESCs. The mESCs were induced to exit pluripotency in medium without Lif for 2 days and then switched to 2i+Lif medium for 5 days. The resultant colonies were fixed and stained for AP. In C and D, the colony morphology and AP intensity were evaluated through microscopy. 100-200 colonies were examined each time to determine the percentage of undifferentiated colonies. The results represent the means (± SD) of three independent experiments. E. Western blotting of pluripotency factors during EB formation. The following figure supplement is available for Figure 2: Figure supplement 1. *Ago2* mRNA is a target of miR-182 and miRNA-183 in mESCs.

Interestingly, *miR-182Δ, miR-183Δ*, and *miR-182Δ/miR-183Δ* mESCs displayed defects in self-renewal (Figure 2C), as determined by the colony formation assay in the 15%FBS+Lif medium, where differentiation was not blocked by the two inhibitors in the 2i+lif medium. Moreover, these miRNA knockout mESCs had accelerated differentiation, as revealed by the exit pluripotency assay (Figure 2D), which evaluates the rate ESCs exit the pluripotent state (Betschinger et al., 2013), and by the measurement of pluripotency factors through Western blotting on differentiating embryonic bodies (Figure 2E). These cellular phenotypes suggest that miR-182/miR-183-mediated regulation of *Ago2* is important to mESCs.

### miR-182/miR-183-mediated repression of *Ago2* is required for maintaining pluripotency

A caveat in interpreting results from miRNA knockout and over-expression experiments is the pleiotropic effects. Because each miRNA can regulate hundreds of mRNAs, when a miRNA is knocked out or over-expressed, hundreds of miRNA:mRNA interactions are altered, making it difficult to determine whether a specific miRNA:mRNA interaction contributes to the phenotypical changes.

To address this issue and specifically examine the functional significance of miR-182/miR-183-mediated regulation of *Ago2* in mESCs, we mutated the miR-182/miR-183 binding sites in the 3’UTR of *Ago2* mRNA via CRISPR/Cas9-mediated genome editing (Figure 3A&B). Two observations indicated that the mutations disrupted the interaction between *Ago2* mRNA and miR-182/miR-183. First, similar to the miRNA knockout mESCs (Figure 2B), Ago2 increased in the 3’UTR mutant mESCs (Figure 3C). Second, in contrast to the results in the WT mESCs (Figure 2 – Figure supplement 1C), over-expression of either miR-182 or miR-183 in the 3’UTR mutant mESCs did not decrease Ago2 (Figure 3 – Figure supplement 1A&B). Notably, the mutations in the 3’UTR of *Ago2* mRNA did not up-regulate Ago2 production *in cis*, because in the *miR-182Δ/miR-183Δ* mESCs, these mutations did not increase Ago2 (Figure 3C), indicating the increased Ago2 from these mutations in the WT mESCs was due to miR-182/miR-183. Moreover, the 3’UTR mutations did not significantly alter the miR-182/miR-183 levels in mESCs (Figure 3 – Figure supplement 1C). Altogether, these observations indicated that the functional significance of miR-182/miR-183-mediated repression of Ago2 could be specifically evaluated in the 3’UTR mutant mESCs.

**Figure 3.**
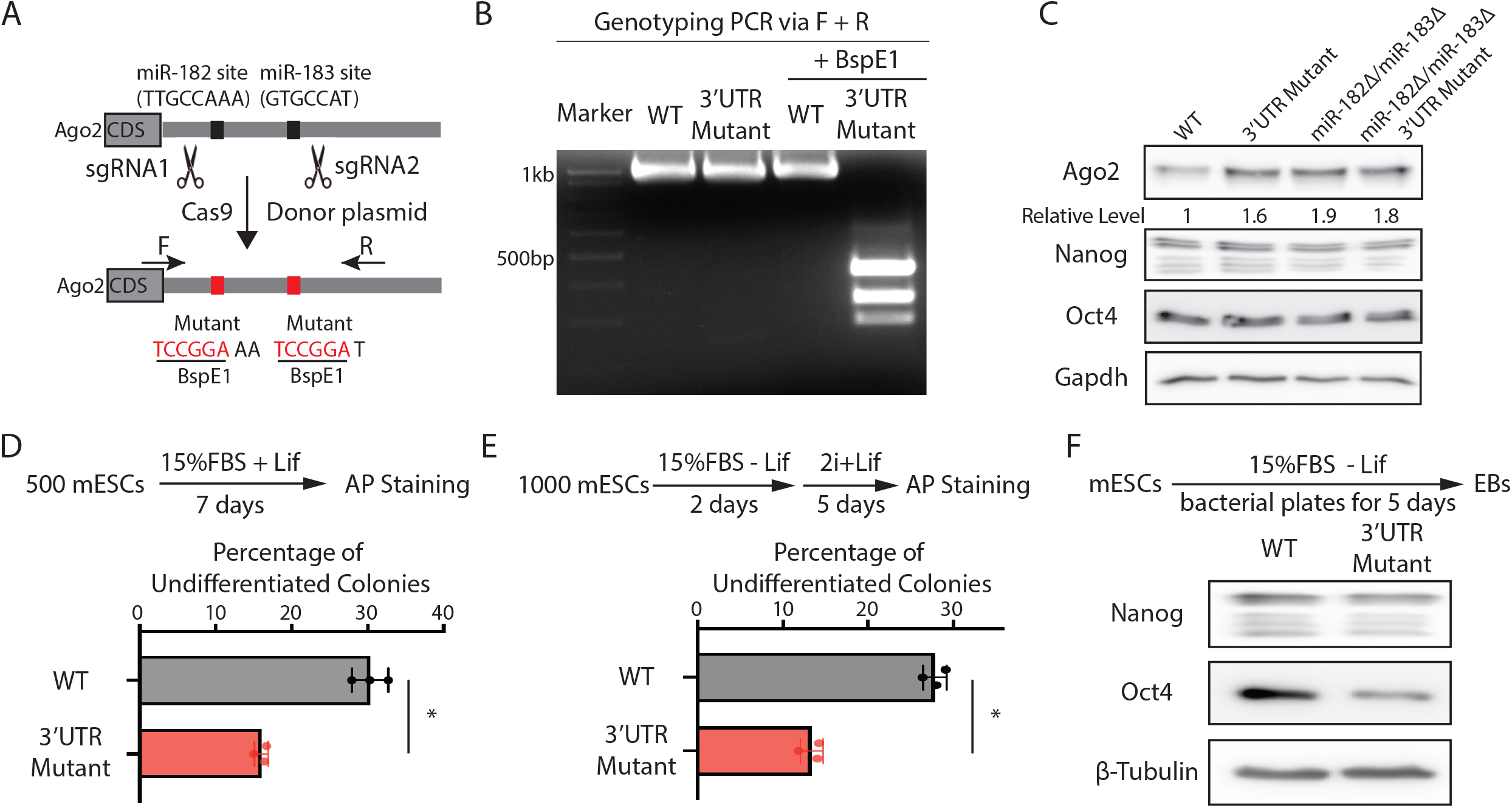
miR-182/miR-183-mediated repression of *Ago2* is required for maintaining pluripotency. A. Mutating miR-182 and miR-183 binding sites in *Ago2* mRNA’s 3’UTR via genome editing. B. Genotyping of the Ago2 3’UTR mutant. The PCR was performed using the oligoes (F and R) indicated in A. C. Western blotting in the WT, *Ago2* 3’UTR mutant, *miR-182Δ/ miR-183Δ*, and *miR-182Δ/ miR-183Δ/Ago2* 3’UTR mutant. D. Colony formation assay for mESCs. E. Exit pluripotency assay for mESCs. In D and E, the colony morphology and AP intensity were evaluated through microscopy. The results represent the means (± SD) of four independent experiments. *p<0.05 by the Student’s t-test. F. Western blotting of pluripotency factors in day 5 EBs. The following figure supplement is available for Figure 3: Figure supplement 1. Inhibition of miR-182- and miRNA-183-mediated regulation of *Ago2* in mESCs.

When subject to the colony formation assay, the 3’UTR mutant mESCs displayed a defect in maintaining undifferentiated colonies (Figure 3D), indicating compromised self-renewal. When differentiation was evaluated by the exit pluripotency assay, the 3’UTR mutant mESCs had an increased differentiation rate (Figure 3E). Consistent with these findings, differentiating embryonic bodies from the 3’UTR mutant mESCs had a lower amount of pluripotency factors (Figure 3F). Collectively, these results indicate that miR-182/miR-183-mediated repression of Ago2 is important for mESC self-renewal and proper differentiation.

### miR-182/miR-183-mediated repression of *Ago2* in mESCs inhibits the let-7 miRNA-mediated differentiation pathway

Two observations lead us to the hypothesis that miR-182/miR-183-mediated repression of *Ago2* in mESCs counteracts the differentiation pathway controlled by the let-7 miRNAs, a conserved miRNA family that promotes stem cell differentiation (Roush and Slack, 2008). First, in *Dgcr8Δ* mESCs, where endogenous miRNAs’ biogenesis is blocked, ectopic expression of miR-183 inhibits the stem cell differentiation triggered by exogenous let-7 miRNA (Wang *et al*., 2017). Second, our recent study indicated that increasing Ago2 levels in mESCs results in stemness defects in a let-7-miRNA-dependent manner. This specificity on let-7 miRNAs is because the pro-differentiation let-7 miRNAs are actively transcribed in mESCs, and the increased Ago2 binds and stabilizes the let-7 miRNAs that are otherwise degraded in mESCs, thereby promoting mESCs differentiation (Liu *et al*., 2021).

To test this hypothesis, we examined the expression of let-7 miRNAs. The 3’UTR mutant mESCs had significantly higher let-7 miRNAs than the WT mESCs (Figure 4A). This increase is specific to let-7 miRNAs because non-let-7 miRNAs were not elevated (Figure 4A). Moreover, consistent with our previous observation that increased Ago2 stabilizes mature let-7 miRNAs (Liu *et al*., 2021), the pri-let-7 miRNAs and the pre-let-7 miRNAs were not significantly increased in the 3’UTR mutant mESCs (Figure 4A). To determine whether the increased let-7 miRNAs are responsible for the stemness defects in the 3’UTR mutant mESCs, we inhibited let-7 miRNAs using locked nucleic acid antisense oligonucleotides (LNA) targeting the conserved seed sequence of let-7 miRNAs. When let-7 miRNAs were inhibited, the stemness defects of the 3’UTR mutant mESCs were abolished (Figure 4B), indicating that disruption of miR-182/miR-183-mediated repression of *Ago2* in mESCs activates differentiation through the let-7 miRNA pathway.

**Figure 4.**
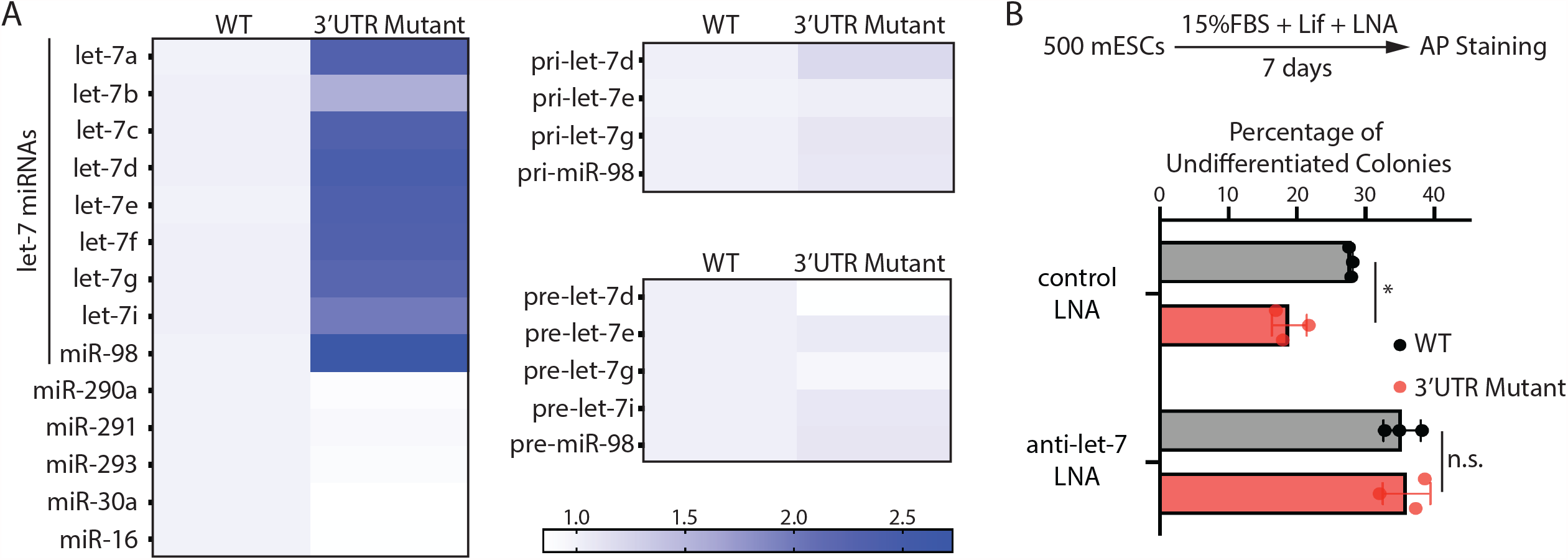
The stemness defects in the 3’UTR mutant mESCs are caused by elevated let-7 miRNAs. A. Relative levels of miRNAs, let-7 pri-miRNAs, and let-7 pre-miRNAs in the WT and the *Ago2* 3’UTR mutant mESCs. For each miRNA, pri-miRNA, and pre-miRNA, the expression level in WT cells was set as 1 for relative comparison. U6 RNA was used for normalization in miRNA and pre-miRNA quantification; 18S rRNA was used for normalization in pri-miRNA quantification. The heatmap was generated from the means of three independent replicates. B. Colony formation assay for WT and the *Ago2* 3’UTR mutant mESCs cultured in the presence of 500 nM anti-let-7 LNA or a control LNA. The results represent three independent experiments. *p<0.05, and n.s. not significant (p>0.05) by the Student’s t-test.

### miR-182/miR-183 and Trim71 function in parallel to repress *Ago2* mRNA in mESCs

Our previous study indicated that *Ago2* mRNA is also repressed by Trim71 in mESCs (Liu *et al*., 2021). Interestingly, the Trim71 binding site in the 3’UTR of *Ago2* mRNA is different from the miR-182/miR-183 binding sites, suggesting that miR-182/miR-183 and Trim71 function in parallel to repress *Ago2* mRNA in mESCs. We performed the following experiments to test this.

At the molecular level, we observed that over-expression of Trim71 still repressed Ago2 in the 3’UTR mutant mESCs (Figure 5A), where miR-182/miR-183-mediated repression is abolished (Figure 3). Moreover, in the 3’UTR mutant mESCs, inhibiting Trim71-mediated repression of Ago2 through deleting the Trim71-binding site in the 3’UTR of *Ago2* mRNA (CLIPΔ) (Liu *et al*., 2021) further increased Ago2 level (Figure 5B, Figure 5 – Figure supplement 1). These results indicate that Trim71 and miR-182/miR-183 independently repress *Ago2* mRNA in mESCs.

**Figure 5.**
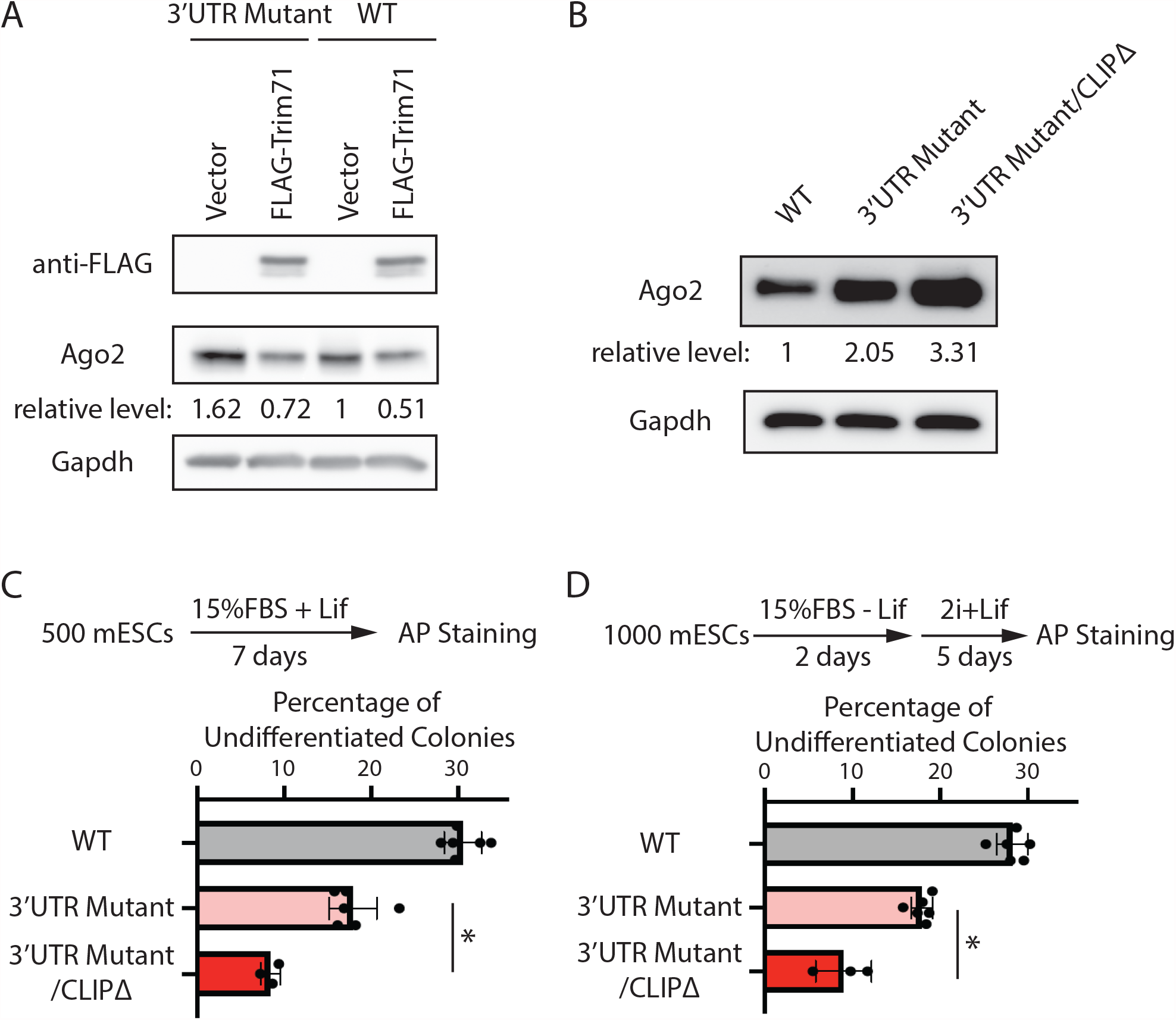
miR-182/miR-183 and Trim71 function in parallel to repress *Ago2* mRNA in mESCs. A. Western blotting in the WT mESCs expressing either a vector or FLAG-Trim71 and in the 3’UTR mutant mESCs expressing either a vector or FLAG-Trim71. B. Western blotting in the WT, 3’UTR mutant, and 3’UTR mutant/CLIPΔ mESCs. In A and B, Gapdh was used for normalization in calculating the relative expression levels. C. Colony formation assay for mESCs. D. Exit pluripotency assay for mESCs. The following figure supplement is available for Figure 5: Figure supplement 1. Generation of the CLIPΔ in the 3’UTR mutant mESCs Supplementary file 1. Antibodies, plasmids, and oligonucleotides used in this study.

At the cell function level, we found that introducing the CLIPΔ in the 3’UTR mutant mESCs further decreased stem cell self-renewal, as determined by the colony formation assay (Figure 5C), and accelerated differentiation, as measured by the exit pluripotency assay (Figure 5D). These observations argue that Trim71 and miR-182/miR-183 function independently in regulating stemness in mESCs through modulating *Ago2* mRNA.

Collectively, these findings indicate that miR-182/miR-183 and Trim71 function in parallel to repress *Ago2* mRNA in mESCs.

## Discussion

Our data reveal that the predominant Ago protein in mESCs, Ago2, is developmentally regulated, with gradually increasing levels when mESCs exit pluripotency. Two miRNAs abundantly expressed in mESCs, miR-182/miR-183, contribute to the repression of *Ago2* in the pluripotent state. This miRNA-mediated regulation of *Ago2* is critical to maintaining stemness. Our findings raise two interesting aspects of miRNAs in stem cell biology.

First, since Ago2 is the predominant Ago protein in mESCs, the Ago2 expression pattern during mESCs’ transition from self-renewal to differentiation argues that although certain individual miRNAs may be required for pluripotency (e.g., miR-182/miR-183), the global miRNA activity is suppressed in the pluripotent state and induced when mESCs initiate differentiation. Consistent with this notion, knocking out key components in global miRNA biogenesis, such as Dgcr8 (Wang et al., 2007), Dicer (Kanellopoulou et al., 2005; Murchison et al., 2005), or Ago2 in the miRISC (Liu *et al*., 2021), does not negatively affect mESCs self-renewal. However, differentiation in all these mutant mESCs is severely compromised. Thus, at the global level, miRNAs may play more important roles in mESC differentiation.

Second, previous studies indicate that the two components of the miRISC, the Ago protein and its associated miRNA, mutually regulate each other. In the absence of miRNAs, the Ago protein is destabilized (Martinez and Gregory, 2013; Smibert et al., 2013), while miRNAs are also unstable if they are not associated with Ago proteins (Winter and Diederichs, 2011). Thus, the effective miRNA activity depends on the limiting component in the miRISC. Our previous studies indicated that the conserved pro-differentiation let-7 miRNAs are sensitive to Ago2 levels because an increase of Ago2 results in specific stabilization of let-7 miRNAs that are otherwise degraded (Liu *et al*., 2021). Thus, for let-7 miRISC, Ago2 is the limiting component in mESCs. Repression of *Ago2* by either miR-182/miR-183 as we characterized here or Trim71 as we identified previously (Liu *et al*., 2021) limits the effective let-7 miRISCs. These two repression mechanisms on *Ago2* mRNA function in parallel to inhibit let-7-mediated differentiation and maintain pluripotency of mESCs. We speculate that similar mechanisms of regulating miRISCs by RNA-binding proteins and microRNAs may exist in other developmental processes. Moreover, Ago2 is dysregulated under many pathological conditions, such as cancer (Adams et al., 2014). Thus, regulating miRISCs through modulating Ago2 levels may also contribute to pathogenesis.

## Materials and Methods

All the antibodies, plasmids, and oligonucleotides used in this study are listed in supplementary file 1.

### CRISPR/Cas9-mediated genome editing in mESCs

To generate the FLAG-Ago1, FLAG-Ago2 mESCs or *Ago2* 3’UTR Mutant mESCs, cells were co-transfected with 2 µg of pWH464 (pSpCas9(BB)−2A-GFP (pX458)) expressing the corresponding targeting sgRNA and 1 µg of the corresponding donor oligo or plasmid using the Fugene6 (Promega). To generate *miR-182Δ* and *miR-183Δ* mESCs, cells were transfected with 2 µg of pWH464 expressing a pair of sgRNAs targeting pri-miR182 or pri-miR183. The transfected cells were subject to single cell sorting and the resulting clones were subject to genotyping to identify the correct clones.

### qRT-PCR

For pri-miRNA quantification, reverse transcription was performed using random hexamers and Superscript2 reverse transcriptase. pre-miRNA and miRNA quantifications were using the Takara’s Mir-X miRNA quantification method. qPCR was performed in triplicate for each sample using the SsoAdvanced Universal SYBR Green Supermix (Bio-Rad) on a CFX96™ real-time PCR detection system (Bio-Rad).

### Western blotting

Proteins were harvested in RIPA buffer (10 mM Tris-HCl pH 8.0, 140 mM NaCl, 1 mM EDTA, 0.5 mM EGTA, 1% Triton X-100, 0.1% sodium deoxycholate, 0.1% SDS, and protease inhibitor cocktail) and quantified with a BCA assay kit (ThermoFisher). Equal amounts of protein samples were resolved by SDS-PAGE gel, and then transferred to PVDF membranes. Western blotting was performed using a BlotCycler (Precision Biosystems) with the corresponding primary and secondary antibodies. The membranes were then treated with the Western ECL substrate (Bio-Rad), and the resulting signal was detected using an ImageQuant LAS 500 instrument (GE Healthcare).

### Colony formation assay and exit pluripotency assay

For colony formation assay, 500 cells were plated on a 12-well plate in 2i+Lif media or Lif media (DMEM/F12 supplemented with 15% FBS, 1 × penicillin/streptomycin, 0.1 mM Non-Essential Amino Acids, 2 mM L-glutamine, and 0.1 mM 2-mercaptoethanol, and 1000 U/ml Lif). For exit from pluripotency assay, 1000 cells were plated on a gelatin-coated 6-well plate in differentiation media (DMEM/F12 supplemented with 15% FBS, 1 × penicillin/streptomycin, 0.1 mM Non-Essential Amino Acids, 2 mM L-glutamine, and 0.1 mM 2-mercaptoethanol) for 2 days, then cultured in 2i+Lif media for another 5 days. Colonies were stained using AP staining Kit and grouped by differentiation status 6-7 days after plating.

### Embryoid body formation

For differentiation via embryoid body formation, 3 × 10^6^ cells were plated per 10 cm bacterial grade Petri dish and maintained on a horizontal rotator with a rotating speed of 30 rpm in differentiation media. The resultant EBs were harvested at the indicated time points.

### RNA antisense purification

mESCs were crosslinked with 0.1% formaldehyde for 5min at room temperature, and the crosslinking reaction was quenched by adding 1/20 volume of 2.5M glycine and incubating the mESCs at room temperature for 10min on a rotating platform. The cells were then harvested and lysed in cell lysis buffer (50mM Tris-HCl pH7.4, 150mM NaCl, 5mM EDTA, 10% glycerol, 1% Tween-20, with freshly added proteinase inhibitors). The cell lysate was cleared by centrifugation at 20,000g for 10min at 4°C. The resulting supernatant was used for RNA antisense purification. 5mg lysate in 500ul lysis buffer was used for each purification. Specifically, a set of 5’-end biotinylated anti-sense DNA oligoes and 5ul RNase inhibitor (NEB) were added to the lysate, resulting in a final concentration of 0.1uM for each oligo. The lysate was incubated at room temperature for 1 hour on a rotating platform. Then 100ul Dynabeads MyOne Streptavidin C1 (Invitrogen) was added and the lysate further incubated for 30min at room temperature on a rotating platform. The magnetic beads were isolated through a magnetic stand and then subject to 4 washes, with each wash in 500ul high salt wash buffer (5XPBS, 0.5% sodium deoxycholate, 1% Triton X-100). The washed beads were resuspended in 100ul DNaseI digestion mix (1X DNase1 digestion buffer with 5ul DNase1 (NEB)) and incubated at 37°C for 20min, followed by adding 350ul LET-SDS buffer (25mM Tris-HCl pH8.0, 100mM LiCl, 20mM EDTA pH8.0, 1%SDS) and 50ul proteinase K (20mg/ml, ThermoFisher). The beads were then incubated on a thermomixer at 55°C 1000rpm for 2 hours. The RNA was isolated through phenol extraction and isopropanol precipitation with glycoblue (Ambion) as a coprecipitant.

## Acknowledgments

We thank Dr. Xiaoli Chen for his assistance with microRNA prediction. This work is supported by Mayo Foundation for Medical Education and Research.

## Supplementary Information

**Figure 1 – Figure supplement 1.**
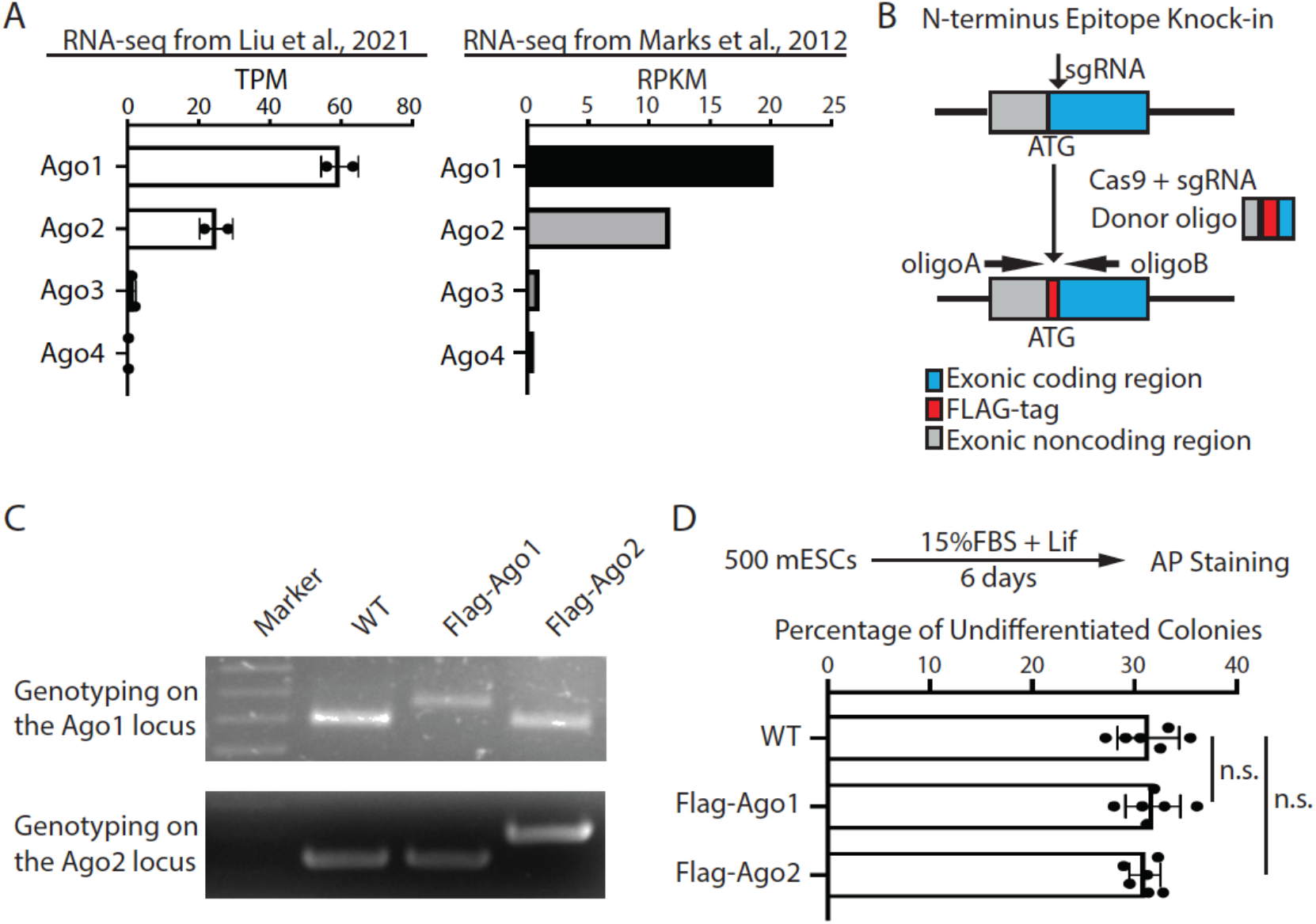
Expression of Argonaute proteins in mESCs. A. mRNA expression levels of *Ago1, Ago2, Ago3*, and *Ago4* in mESCs. TPM: transcripts per kilobase million mapped reads; RPKM: reads per kilobase of transcript per million mapped reads. B. Workflow for N-terminus knock-in the Flag-tag to the endogenous Ago1 and Ago2 loci in mESCs. C. Genotyping of the Flag-Ago1 and the Flag-Ago2 mESCs. D. Colony formation assay for the WT, Flag-Ago1, and Flag-Ago2 mESCs. The results represent the means (± SD) of six independent experiments, and in each experiment, 100–200 colonies were evaluated for each type of cells. n.s. not significant (p>0.05) by the Student’s t-test.

**Figure 1 – Figure supplement 2.**
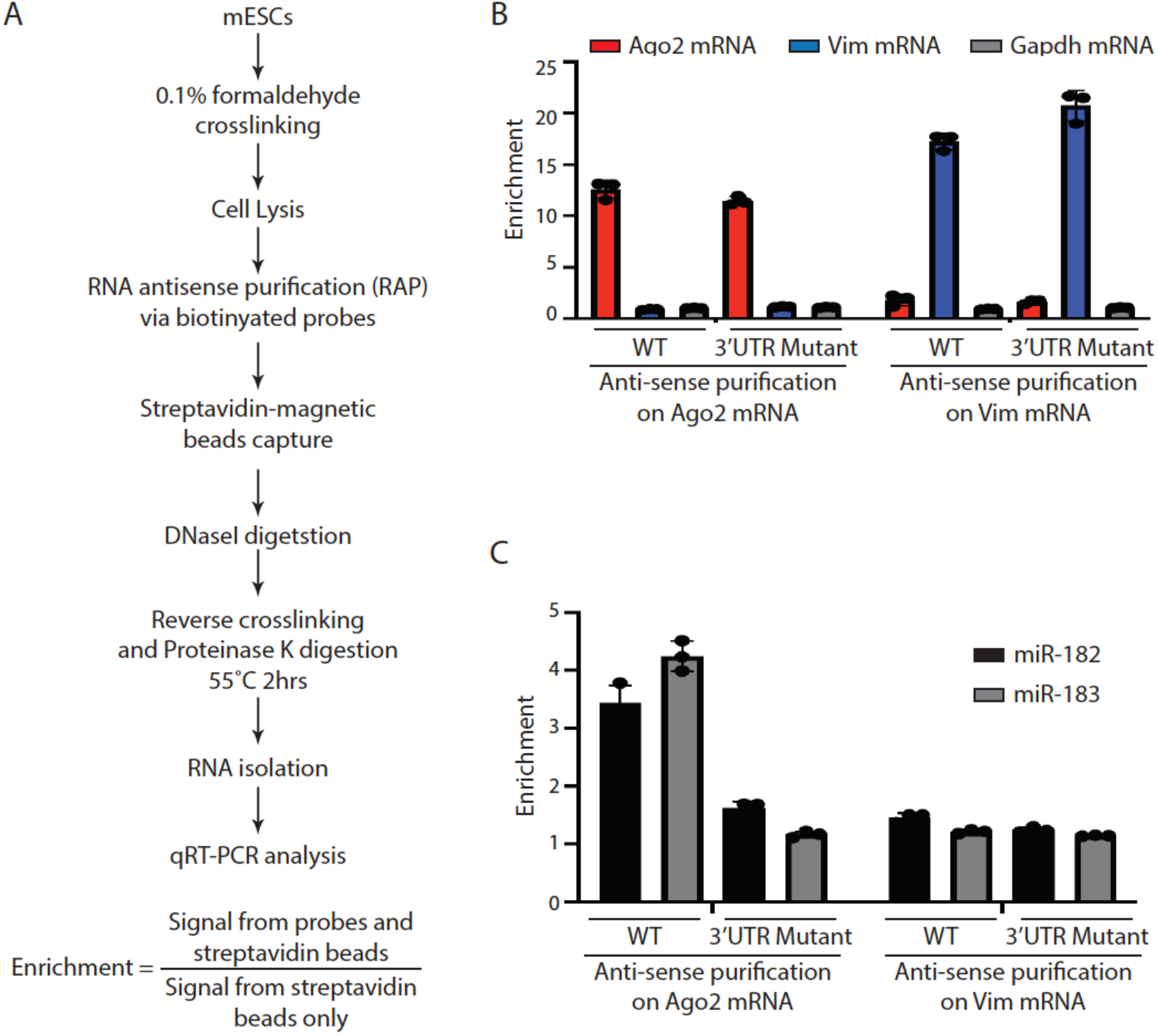
miR-182 and miR-183 are specifically associated with *Ago2* mRNA in mESCs. A. Outline of the modified RNA anti-sense purification procedures. B. qRT-PCR quantification of mRNAs isolated from the RNA anti-sense purification in the WT and the 3’UTR mutant mESCs. C. qRT-PCR quantification of microRNAs isolated from the RNA anti-sense purification in the WT and the 3’UTR mutant mESCs. The results in B and C represent the means (± SD) of three independent experiments.

**Figure 2 – Figure supplement 1.**
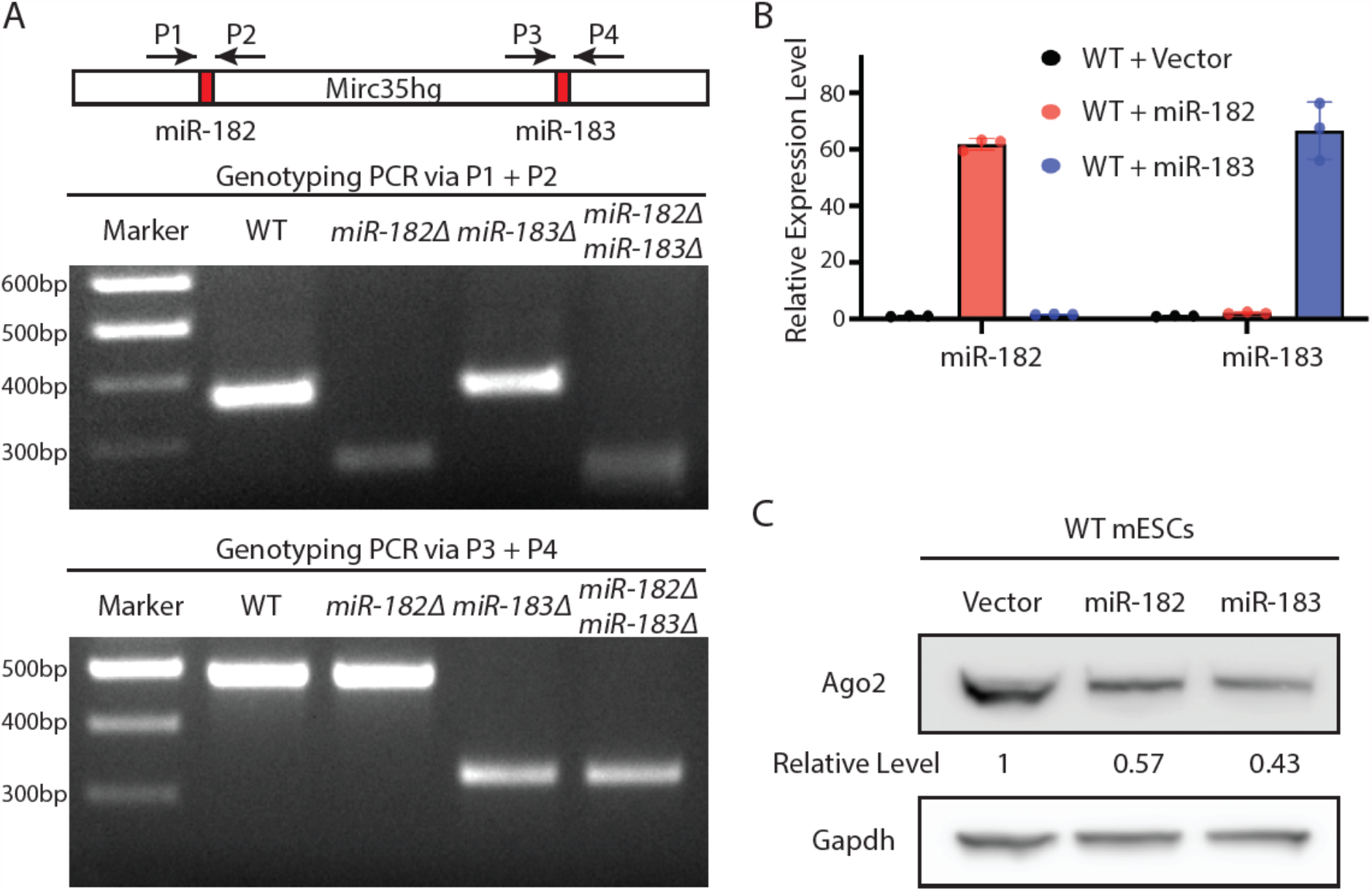
*Ago2* mRNA is a target of miR-182 and miRNA-183 in mESCs. A. Genotyping for the *miR-182Δ, miR-183Δ*, and *miR-182Δ/ miR-183Δ* mESCs. B. qRT-PCR on miR-182 and miR-183 in mESCs with either miR-182 or miR-183 overexpression. The results represent the means (± SD) of three independent experiments, and U6 RNA was used for normalization. C. Western blotting in the WT mESCs overexpressing either miR-182 or miR-183.

**Figure 3 – Figure supplement 1.**
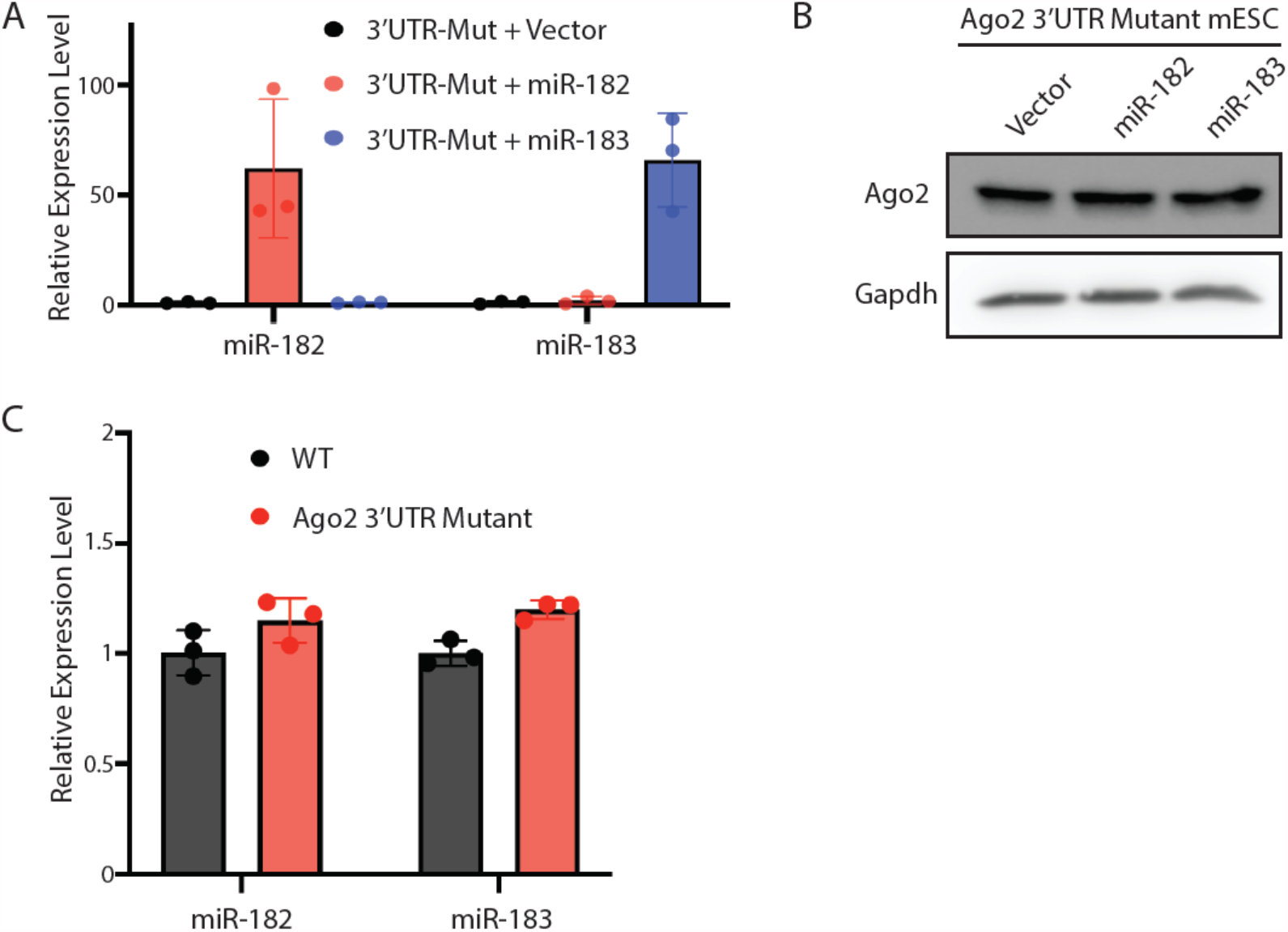
Inhibition of miR-182- and miRNA-183-mediated regulation of *Ago2* in mESCs. A. qRT-PCR on miR-182 and miR-183 in the *Ago2* 3’UTR mutant mESCs with either miR-182 or miR-183 overexpression. B. Western blotting in the *Ago2* 3’UTR mutant mESCs overexpressing either miR-182 or miR-183. C. qRT-PCR on the endogenous miR-182 and miR-183 in the WT and the *Ago2* 3’UTR mutant mESCs. The results of (A) and (C) represent the means (± SD) of three independent experiments, and U6 RNA was used for normalization.

**Figure 5 – Figure supplement 1.**
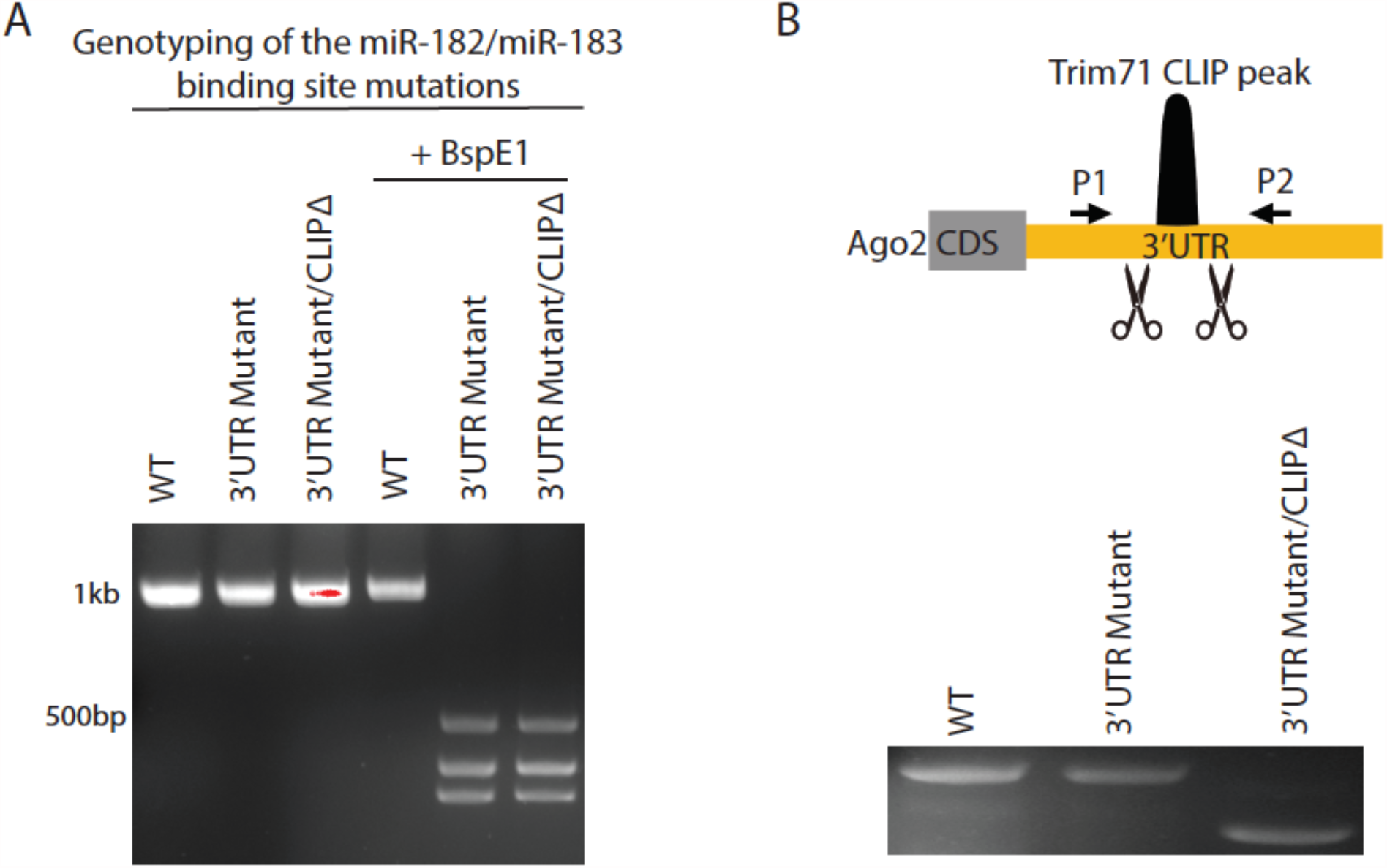
Generation of the CLIPΔ in the 3’UTR mutant mESCs. A. Genotyping for the 3’UTR mutant mESCs. The genotyping primers were indicated in Figure 3A. B. Genotyping for the CLIPΔ.

